# Targeted brain tumor radiotherapy using an Auger emitter

**DOI:** 10.1101/649764

**Authors:** Giacomo Pirovano, Stephen A. Jannetti, Lukas M. Carter, Ahmad Sadique, Susanne Kossatz, Navjot Guru, Paula Demetrio De Souza Franca, Masatomo Maeda, Brian M. Zeglis, Jason S. Lewis, John L. Humm, Thomas Reiner

**Affiliations:** Department of Radiology, Memorial Sloan Kettering Cancer Center, 1275 York Avenue, New York, NY, 10065, USA; Department of Biochemistry, Hunter College - The City University of New York (CUNY), New York, NY, 10028, USA; Ph.D. Program in Biochemistry, The Graduate Center, CUNY, New York, NY, 10016, USA; Department of Chemistry, Hunter College of the City University of New York, New York, NY, 10028, USA; Department of Radiology, Weill Cornell Medical College, New York, NY, 10065, USA; Ph.D. Program in Chemistry, the Graduate Center of the City University of New York, New York, NY, 10016, USA; Molecular Pharmacology Program, MSK, New York, NY, 10065, USA; Department of Pharmacology, Weill-Cornell Medical College, New York, NY, 10065, USA; Department of Medical Physics, Memorial Sloan Kettering Cancer Center, New York, NY, 10065, USA; Chemical Biology Program, Memorial Sloan Kettering Cancer Center, New York City, NY 10065, USA

## Abstract

Glioblastoma multiforme is a highly aggressive form of brain cancer whose location, tendency to infiltrate healthy surrounding tissue, and heterogeneity significantly limit survival, with scant progress having been made in recent decades. ^123^I-MAPi (Iodine-123 Meitner-Auger PARP1 inhibitor) is a precise therapeutic tool composed of a PARP1 inhibitor radiolabeled with an Auger- and gamma-emitting iodine isotope. Here, the PARP inhibitor, which binds to the DNA repair enzyme PARP1, specifically targets cancer cells, sparing healthy tissue, and carries a radioactive payload within reach of the cancer cells’ DNA. The high relative biological efficacy of Auger electrons within their short range of action is leveraged to inflict DNA damage and cell death with high precision. The gamma ray emission of ^123^I-MAPi allows for the imaging of tumor progression and therapy response, and for patient dosimetry calculation. Here we demonstrated the efficacy and specificity of this small molecule radiotheranostic in a complex preclinical model. *In vitro* and *in vivo* studies demonstrate high tumor uptake and a prolonged survival in mice treated with ^123^I-MAPi when compared to vehicle controls. Different methods of drug delivery were investigated to develop this technology for clinical applications, including convection enhanced delivery (CED) and intrathecal injection. Taken together, these results represent the first full characterization of an Auger-emitting PARP inhibitor, demonstrate a survival benefit in mouse models of GBM, and confirm the high potential of ^123^I-MAPi for clinical translation.

**One Sentence Summary:** A novel PARP1-targeted Auger radiotherapeutic shows translational potential as a theranostic tool for imaging and killing cancer cells, resulting in tumor delineation and prolonged survival in a glioblastoma model.

---

Glioblastoma multiforme (GBM) is one of the deadliest forms of solid tumors, with a 5-year survival rate as low as 5% (*1*). Clinical intervention typically consists of maximal safe surgical resection followed by adjuvant chemo-radiotherapy. This therapeutic regimen, unfortunately, imparts only limited improvements to survival (*2*). Additionally, the GBM molecular heterogeneity represents a robust challenge in need of better imaging tools that would allow for the monitoring of therapy response and lead to better and more personalized therapeutic plans. Furthermore, the presence of the blood-brain barrier (BBB) biochemically limits the pharmacokinetics of many GBM drug candidates.

A novel approach for treating GBM is therefore urgently needed, one that would address both pharmacodynamic as well as pharmacokinetic hurdles (*3, 4*). Promising delivery strategies to overcome these problems in the clinics include convection enhanced delivery (CED) (*5, 6*) and intrathecal injection (*7, 8*).

A known molecular biomarker for most tumors, including GBM, is poly(ADP-ribose) Polymerase 1 (PARP-1) (*9–15*). PARP-1 is recruited to the nucleus of cancer cells and binds DNA as a single-strand break (SSB) repair enzyme (*16*). This central role has been successfully leveraged for the development of various PARP inhibitors, both as a monotherapy and in combination with other therapeutics (*17, 18*). Modified PARP inhibitors have also been widely used for tumor detection and imaging due to their cancer specificity and their high target-to-background contrast (*14, 15, 19–25*); more recently, they have found theranostic applications (*26*).

Here, we focus on a less used type of radioactive emission: Auger radiation. Auger emitters are an extremely potent radioactive source for targeted radiotherapy, characterized by their greater linear energy transfer, incredibly short range, and ability to cause a complex, lethal DNA damage as compared to traditional X-rays or β-particles (*27–30*). Previous attempts to use Auger emitters as cancer therapies have not been successful, due to the limited range of the radiation emitted and the difficulty of reliably delivering the lethal electrons close enough to the DNA target (< 100 Å) (*31–33*).

In this study, we developed and characterized ^123^I-MAPi, the first Auger-based theranostic PARP inhibitor able to directly deliver its lethal payload within a 50-angstrom distance of the DNA of GBM cancer cells (fig. 1A). This distance is within the Auger radius of action, resulting in an effective preclinical cancer treatment drug and leading to improved survival in a preclinical GBM model. We used the 159 keV gamma ray to image tumor progression and calculate dosimetry and treatment efficacy using SPECT imaging.

**Figure 1.**
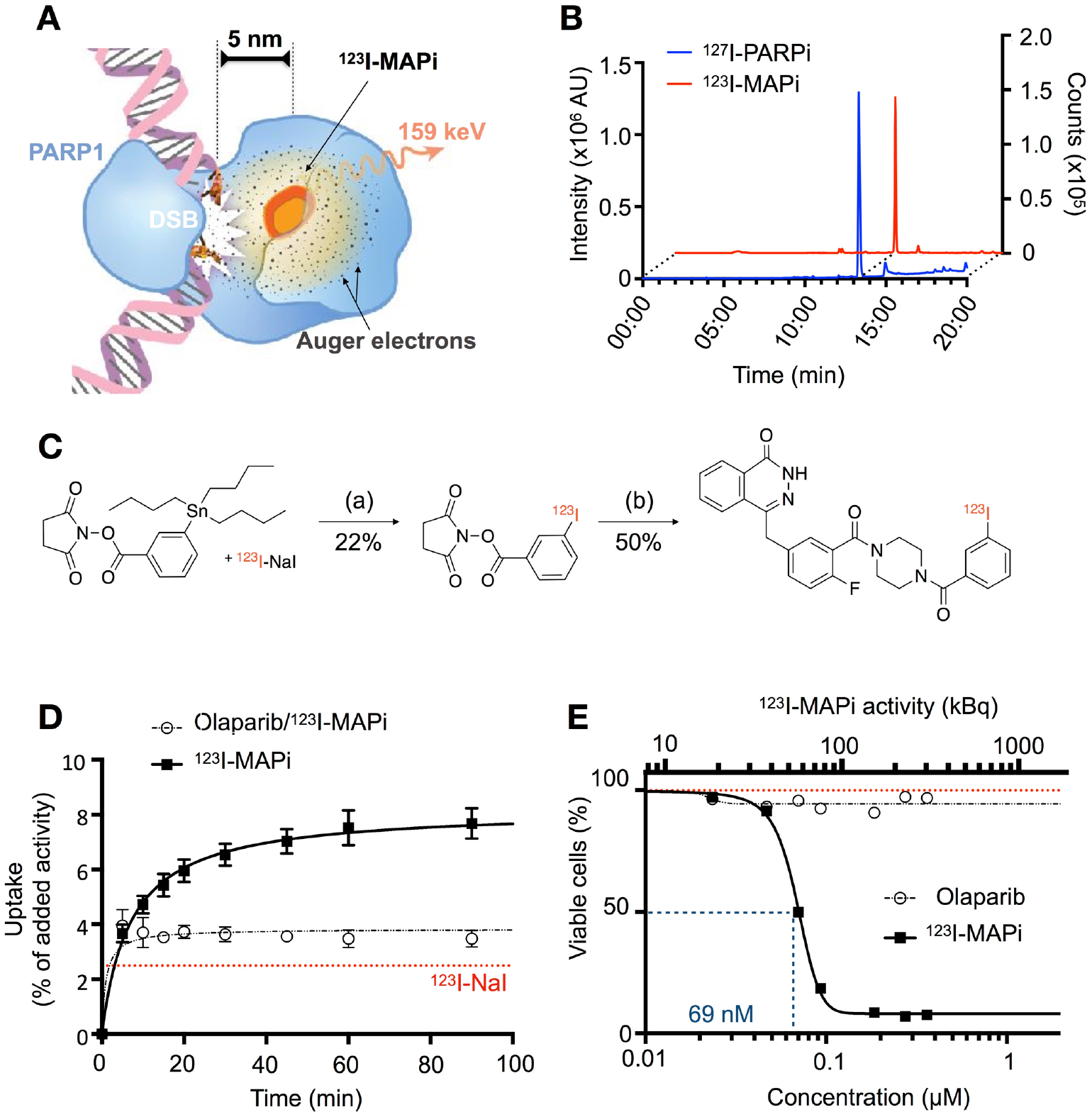
^123^I-MAPi Synthesis, binding, and efficacy *in vitro*. **A.** ^123^I-MAPi binding to PARP1 can deliver lethal Auger radiation within ~100 Å, enough to affect DNA in cancer cells when bound. **B.** RP-HPLC coelution of ^123^I-MAPi with nonradioactive I-127 analog. **C.** Synthetic scheme of ^123^I-MAPi, (a) 0.1 M NaOH, Chloramine T, AcOH, MeOH 20 m, RT; (b) 4-(4-fluoro-3-(piperazine-1-carbonyl)benzyl)phthalazin-1(2*H*)-one, HBTU, DMAP, 2,6-Lutidine, DMSO, ACN, 2 h, 65 °C. **D.** Cellular uptake of ^123^I-MAPi in U251 GBM cell line. Blocking performed with 100-fold incubation of Olaparib prior to treatment. Sodium Iodine-123 represented by red line. Michaelis-Menten curve fitting. **E.** Alamar Blue assay comparing ^123^I-MAPi efficacy with Olaparib at equal molar concentrations. ^123^I-MAPi EC_50_ = 69 nM. Nonlinear fit four parameters variable slope.

Taken together, these results illustrate the tremendous potential of ^123^I-MAPi as an Auger-emitting PARP inhibitor and a theranostic agent for GBM treatment.

## RESULTS

### Synthesis of ^123^I-MAPi and in vitro validation

We previously showed that it is possible to successfully conjugate a PARP inhibitor with radioiodine without altering the high affinity to the target with an IC_50_ in the nanomolar range (11 ±3 nM) and a logPCHI of 2.3 (*34*). ^123^I-MAPi is a novel, previously unreported isotopologue, and was synthesized with a final molar activity of 3.93 ± 0.10 GBq/μmol. Radiochemical purity was 99.1 ± 0.9% for all prepared compounds (fig. 1B).

Pharmacological properties determined with ^127^I-PARPi suggest that ^123^I-MAPi (fig. 1B, S1B), retains the same properties as ^131^I-PARPi, which have been shown to be similar to the FDA-approved PARP inhibitor Olaparib (*26, 34*).

*In vitro* internalization was tested on U251 cells expressing PARP1 (fig. 1D, S1A). ^123^I-MAPi (37 kBq/well) was added to adherent cells in monolayer and uptake was calculated by measuring gamma radiation in cell lysates at different time points. We confirmed rapid cellular internalization, with 50% of the final total uptake being reached after 5 – 10 minutes postincubation and with an uptake plateau reached at 1 h post-treatment. The final total uptake was ~8% of added activity, V_max_ = 8.2 ± 0.2 % compared to blocked uptake, V_max_ = 3.8 ± 0.1 % (Michaelis-Menten fit, R^2^ = 0.974 and R^2^ = 0.879, respectively). Uptake was blocked with a 100-fold excess dose of Olaparib to show target specificity. Na-^123^I was used as a control and showed significantly lower uptake, V_max_ = 2.5 ± 0.1. The two dominant targets for ^127^I-PARPi are PARP1 and PARP2 (*26*), similar to what has been previously reported for Olaparib and other modified PARP inhibitors (*15*).

We tested the cancer-specific efficacy of ^123^I-MAPi *in vitro* by treating U251 GBM cells with increasing doses (fig. 1E). ^123^I-MAPi proved to be capable of killing cancer cells with a submicromolar EC_50_ (EC_50_ = 68.9 ± 1.1 nM, R^2^ = 0.999). At the same concentrations, Olaparib did not show any effect in terms of cell viability. Comparing ^123^I-MAPi with the previously-published beta-emitting ^131^I-PARPi, we observed a 16-fold increase in EC_50_ potency (EC_50_ of ^131^I-PARPi being 1148 ± 1 nM, fig. S1C). ^123^I-MAPi treatment proved to reduce clonogenicity and viability when compared to ^123^I-NaI (fig. S1D and S1E), suggesting that the observed effect is due to the PARP inhibitor mediated close proximity to the target DNA.

### Biodistribution and toxicity of ^123^I-MAPi in vivo

TS543 patient-derived glioblastoma stem cells were used to grow tumors in athymic nude mice in order to investigate the biodistribution of ^123^I-MAPi. Tumors were grown subcutaneously on the animals’ right shoulder. Mice were then randomized and divided into two cohorts (n = 3/group) one of which was used for blocking. Blocking was performed systemically with an intravenous injection of Olaparib 1 h prior to ^123^I-MAPi treatment. ^123^I-MAPi was injected intratumorally for this model. SPECT/CT images showed that ^123^I-MAPi tumor uptake was retained at 1 h, 6 h and 18 h post injection in non-blocked animals, but was strongly reduced at 6 h and 18 h in blocked animals (fig. 2A, fig. S2A). Biodistribution at 18 h was also examined and showed higher tumor uptake (33.4 ± 28.0 % ID/g) compared to the tumor of blocked animals (0.4 ± 0.1 % ID/g) (fig. 2B). Tumor-to-muscle ratios in ^123^I-MAPi treated mice versus blocked mice were > 500 and 5, respectively, with less than 1% ID/g in all clearing organs.

**Figure 2.**
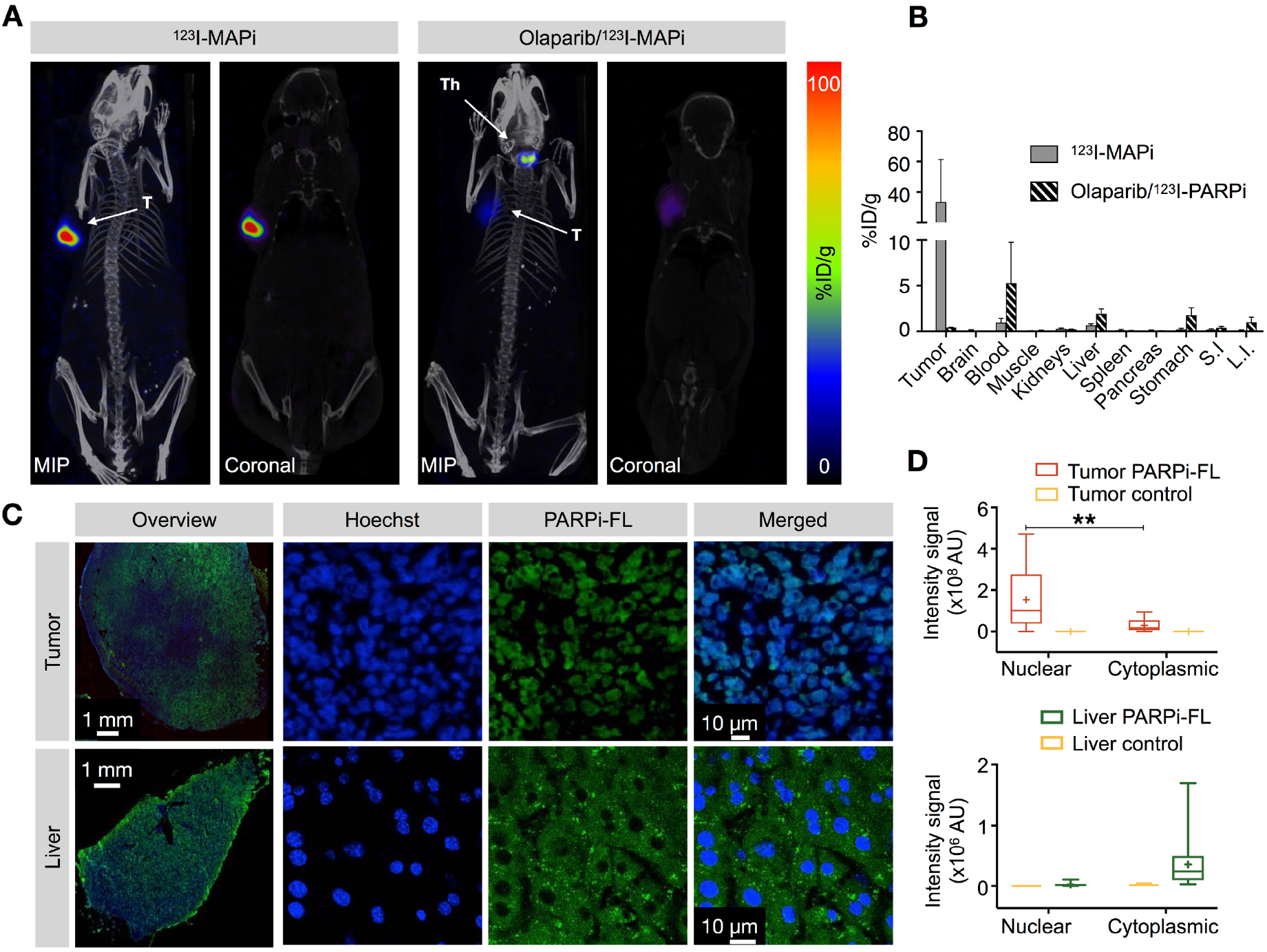
Imaging and biodistribution of ^123^I-MAPi in subcutaneous TS543 mouse model. **A.** Specific tumor uptake of ^123^I-MAPi at 18 h after local injection. Blocking was performed intravenously 1 h before injection. **T** = tumor, **Th** = thyroid. **B.** *Ex vivo* biodistribution of ^123^I-MAPi at 18 h after local injection. **C.** Uptake of PARPi-FL in tumor and liver of TS543 tumor-bearing mice show nuclear uptake for tumor and cytoplasmic uptake for liver tissue. **D.** Quantification of PARPi-FL uptake in the nucleus and cytoplasm of tumor and liver tissue. Tumor control was performed by IV injection of blocking Olaparib dose. Liver control was performed by injection of vehicle. Nuclear vs cytoplasmic uptake was automatically detected using an ImageJ script to detect colocalization of Hoechst (IV injected 5 minutes before extracting the organs) and PARPi-FL. Student t-test, \*\**p*-value < 0.01, \*\*\**p*-value < 0.001.

Auger particles are highly cytotoxic when they can directly interact with the DNA and cause complex damage. However, they are significantly less so in the cellular cytosol, where they are beyond the reach of their DNA target (*27*). As liver clearance is the main route of excretion of ^123^I-MAPi that could cause dose-limiting problems in the clinic, we investigated potential clinical liver failure by observing specific liver toxicity in our preclinical study. We compared tumor and liver accumulation of PARPi-FL, a thoroughly characterized fluorescent analogue of the same PARP1 inhibitor (*14, 15, 20, 34, 35*). PARPi-FL was injected intravenously (50 μg per mouse) 2 h before collecting the animals’ tumor and liver (fig. 2C and fig. S3A). Organs were then sectioned, and tissue sections digitalized. Hoechst (150 μL/mouse of 10 mg/mL) was used for nuclear counterstaining. PARPi-FL accumulated in the nucleus of GBM cells as confirmed by co-localization with Hoechst signal. Livers were collected and imaged with an inverted confocal microscope. PARPi-FL accumulation was observed in the cytoplasm of liver cells (fig. 2D), suggesting that liver cells are protected from Auger-toxicity during hepatobiliary clearance, due to the large distance of the Auger emitter from the DNA (fig. S3B).

### Imaging of an orthotopic GBM model using ^123^I-MAPi

To test ^123^I-MAPi in a more realistic model of GBM, we orthotopically implanted TS543 cells into the right brain hemisphere. This GBM model proved to be very consistent in growth and take rate, as we monitored by MRI imaging of the head. 50,000 cells were injected at Week 0, which resulted in rapid disease progression leading to animal death at Week 7 (fig. 3A). Based on this data we decided to deliver ^123^I-MAPi treatment at Week 3. ^123^I-MAPi was injected intratumorally using the same stereotactic coordinates as for tumor implantation. Animals were then imaged at 1 h and 18 h post-treatment. Full body SPECT/CT images were acquired, showing retention of ^123^I-MAPi in the tumor at 18 hours post-injection (fig. 3B and fig. S4A).

**Figure 3.**
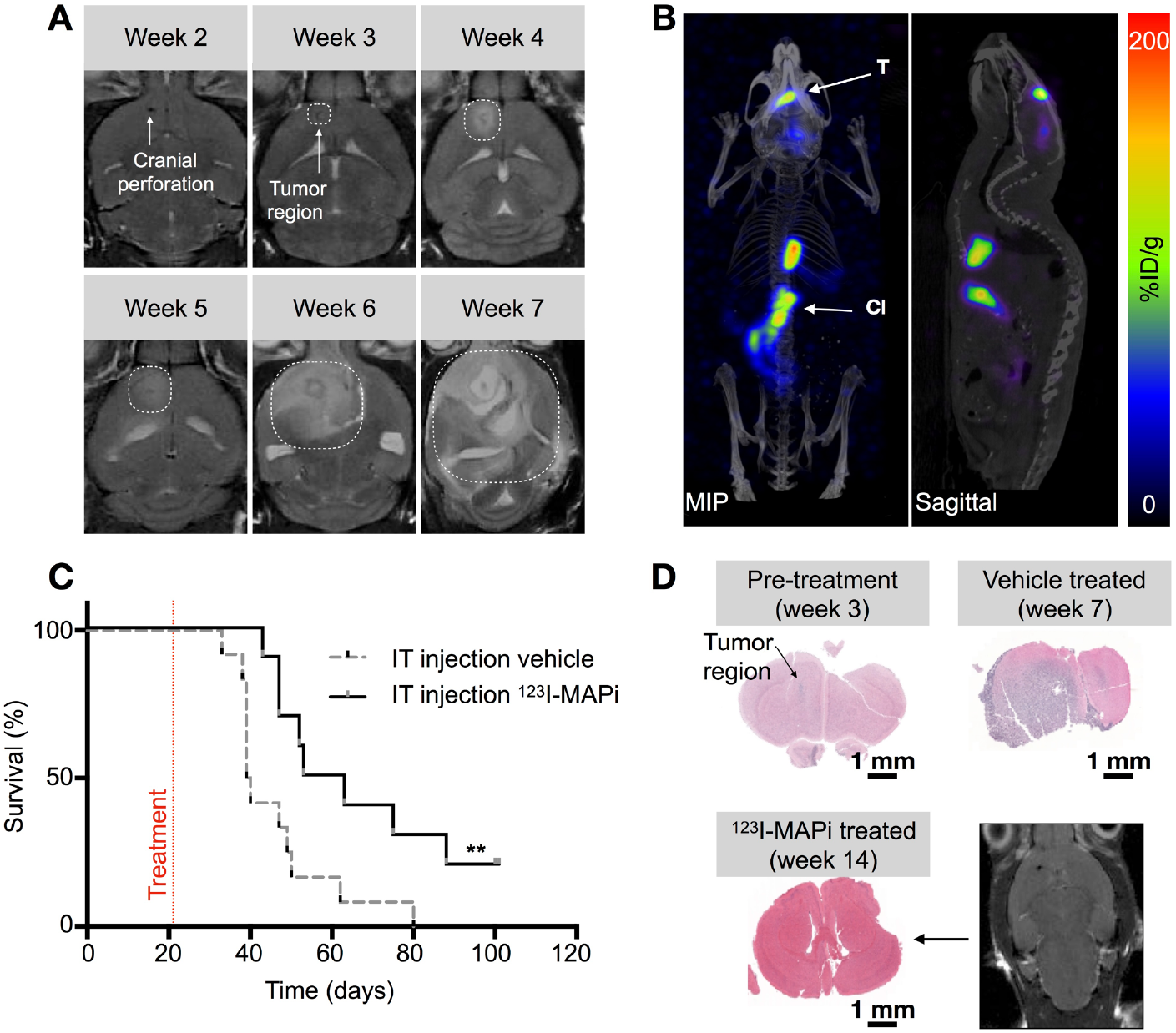
TS543 glioblastoma stem-cell mouse model and ^123^I-MAPi efficacy. **A.** MRI monitoring of TS543 xenograft disease progression. **B.** SPECT/CT imaging of GBM with ^123^I-MAPi single injection. Images were taken at 18 h after local injection. **T** = tumor, **Cl** = clearing organs. **C.** Kaplan-Meier curve of mice injected with ^123^I-MAPi (local single injection 370 kBq - 1.11 MBq, n = 10) compared to vehicle injection (n = 12). Log-Rank (Mantel-Cox) test, \*\**p*-value < 0.01. **D.** Cytology staining of untreated mice at Weeks 3 and 7 after tumor implantation and cytology staining and MRI imaging of treated mouse brain at 14 weeks post implantation.

### Therapeutic efficacy of ^123^I-MAPi in a GBM mouse model

We monitored the clinical potential utility of ^123^I-MAPi in treating GBM in the above described orthotopic mouse model. ^123^I-MAPi-treated mice were then monitored daily, using Day 98 (the end of Week 14), measured from the day of xenografting, as the study endpoint. We injected an intratumoral single dose (0.37 – 1.11 MBq) of ^123^I-MAPi for the treated cohort (n = 10) and the same volume of vehicle for the control cohort (n =12). Survival data confirmed an improved survival for the ^123^I-MAPi treatment cohort, with a median survival of 58 days as opposed to 40 days observed for the control cohort. Log-rank curve comparison showed a significant difference, with p-value = 0.009 (fig. 3C, table 1). Animals brains were imaged *ex vivo* with H&E staining and *in vivo* with MRI showing a diffuse presence of cancer cells in vehicle treated mice as opposed to the treatment (fig. 3D).

**Table 1.** Survival of single dose treatment with ^123^I-MAPi.

|  | **n** | **Median surv. (days)** | ***p*-value<sup>†</sup>** |
| --- | --- | --- | --- |
| **IT injection vehicle** | 12 | 50 |  |
| **IT injection $^{123}\text{I}$-MAPI** | 10 | 58 | **\*\* 0.0094** |

<sup>†</sup>Log-Rank (Mantel-Cox) test.

### Improved delivery of ^123^I-MAPi for clinical translation

In clinics, to improve biodistribution toward a more effective targeted therapy, the radiopharmaceutical can be administered directly into the tumor compartment. For brain tumors this means intra-tumoral injection, intra-thecal injection, or convection enhanced delivery (CED), an approach that elevates the injection pressure so as to impel the agent across the BBB. To allow for clinical translation of ^123^I-MAPi, we first built a preclinical model of CED. We implanted an ALZET^®^ osmotic delivery pump with a subcutaneous catheter in tumor-bearing mice. This delivered the content of the subcutaneous reservoir at a flow rate of 1 μL/h over the course of approximately 100 h through a cannula connected to the mouse brain (fig. S6A). Dosimetry for the CED experiments (fig. 4A) confirmed accumulation in the tumor with minimal dose to healthy organs not in direct proximity to the implanted pump (fig. S6B, S6C, and S6D). Importantly, while not feasible in a mouse model, normal organ doses would be *significantly* reduced in a corresponding clinical scenario where the radionuclide reservoir would be external and shielded. Subcutaneous reservoirs were filled with 100 μL of ^123^I-MAPi for the treatment cohort (n = 8) and with 100 μL of vehicle for the control cohort (n = 8). Five days post-implantation, we surgically removed the pumps, as per manufacturer instruction, and monitored mice survival. Kaplan-Meier survival plots confirmed the therapeutic efficacy of ^123^I-MAPi: treated mice presented a median survival of 72 d as opposed 48 d in the control cohort, a statistically significant difference of 50% (Log-rank *p*-value = 0.0361, fig. 4B and table 2).

**Figure 4.**
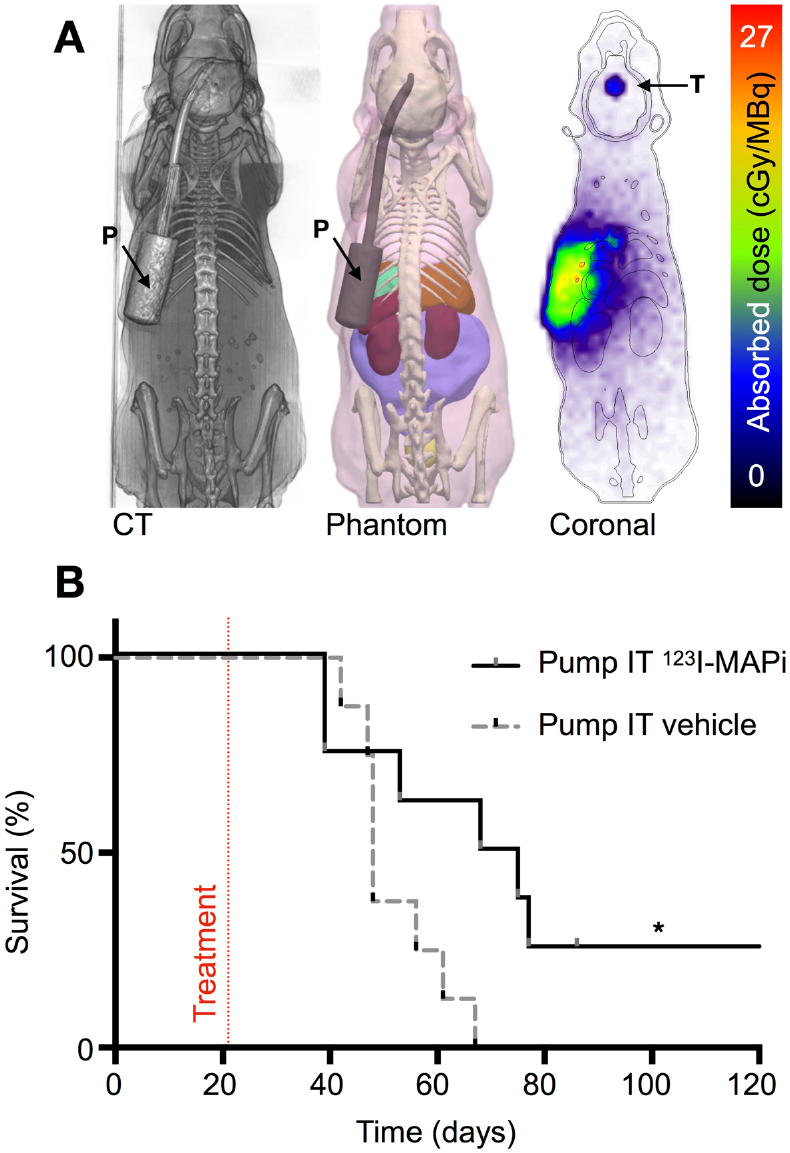
Improved delivery of ^123^I-MAPi with an *in vitro* CED model. **A.** Dosimetry of the subcutaneous pump model showing CT, phantom and Monte Carlo simulation of dose accumulation in the tumor. **B.** Kaplan-Meier survival study of pump implanted mice shows an improvement of survival of ^123^I-MAPi treated mice (n = 8) when compared to control (n = 8). Treatment mice osmotic pumps were loaded with 481 ± 111 kBq. Log-Rank (Mantel-Cox) test, \**p*-value < 0.05.

**Table 2.** Survival of CED dose treatment with ^123^I-MAPi.

|  | **n** | **Median surv. (days)** | ***p*-value<sup>†</sup>** |
| --- | --- | --- | --- |
| **IT injection vehicle** | 8 | 48 |  |
| **IT injection $^{123}\text{I}$-MAPI** | 8 | 72 | **\* 0.0361** |

<sup>†</sup>Log-Rank (Mantel-Cox) test.

### Intrathecal injection of ^123^I-MAPi for a further simplified clinical translation

We then investigated a more broadly used and technically feasible brain drug delivery method for delivering ^123^I-MAPi across the BBB. Tumor-bearing mice were injected with 50 μL of ^123^I-MAPi intrathecally at 3 weeks after orthotopic tumor implantation. SPECT/CT imaging was performed at 1 h post injection in order to visualize ^123^I-MAPi accumulation in the brain (fig. S5A). Imaging confirmed favorable pharmacokinetics and specific tumor uptake, suggesting intrathecal delivery as a feasible technique for our small molecule. Unfortunately, this technique lead to surgery-related high levels of stress in the mice model preventing us from being able to monitor survival.

## Discussion

In the present work, we show the results for the first preclinical characterization of an Auger-emitting theranostic PARP inhibitor. We present a functioning workflow for the synthesis of a stable ^123^I-MAPi compound which proved to be effective *in vitro* at reducing the viability of GBM cell lines. The lethality of ^123^I-MAPi suggests that the DNA is in range of the Auger electrons when it is complexed with PARP1. It was further established that GBM cell death is induced through radiogenic damage rather than PARP inhibition at these tracer levels.

*In vivo* studies with human tumor xenografts models also show ^123^I-MAPi drug treatment efficacy. The 159 keV gamma emission of ^123^I allowed for quantification of drug uptake in tumor-bearing mice by SPECT/CT imaging. The isotope is ideal isotope for a proposed theranostic PARPi clinical agent because of its widespread use in nuclear medicine for thyroid disorders (Na^123^I) and pediatric tumors (^123^I-MIBG). Animal studies employing the nonradioactive drug Olaparib show prolongation of blood pool clearance and blocked tumor uptake, proving ^123^I-MAPi binding specificity.

To illustrate the potential advantages of clinical translation of ^123^I-MAPi, and to better illustrate the impact of Auger therapeutics, we looked at the agent’s potential subcellular biodistribution. Olaparib-based PARP inhibitors are cleared mainly hepatobiliary (*15*) and the liver is therefore a potential organ for Auger-specific radiotoxicity. In preparation for clinical translation of ^123^I-MAPi, we performed studies to investigate liver toxicity. We found near-exclusive extracellular or perinuclear localization of ^123^I-MAPi in liver cells (i.e. the DNA of the liver parenchymal cells is out of range of most of the emitted low-range Auger electrons) as shown with the fluorescent analog PARPi-FL. This is in stark contrast to the observed nuclear accumulation of ^123^I-MAPi in GBM cells (fig. 5B).

To force a more favorable biodistribution toward tumor-targeting in clinical targeted radionuclide studies directed at brain tumors, investigators have proposed intra-tumor injection (*36*), intrathecal injection (*37*) and more recently convection enhanced delivery (CED) to administer the antibody across the BBB and into tumor tissue – e.g. to treat diffuse intrinsic pontine glioma (*38*). The above approaches have shown the ability to significantly improve the ratio of the radiation dose delivered to tumor relative to normal dose-limiting tissues. To emulate these studies in a pre-clinical setting, we adopted an orthotopic GBM model in mice and performed a local injection of ^123^I-MAPi using an intratumoral osmotic pump delivery system. This system allowed for constant and prolonged delivery of the drug directly to the tumor and its microenvironment. Dosimetry estimates based on imaging, tissue specimen counting, and treatment response shown by Kaplan-Meier survival data are suggestive of the high potential clinical impact of this approach.

CEDs and Ommaya reservoirs are emerging as effective chemotherapy and targeted radionuclide delivery methods for patients with previously inaccessible metastatic disease. We have developed a method to perform CED injection of novel radiopharmaceuticals in preclinical models of cancer. We also performed intrathecal injection which would allow tumor imaging and therapy in clinical settings. This technique is routinely performed in most hospitals for chemotherapy drug delivery and allows clinicians to access the brain without opening the patient’s skull. SPECT/CT imaging can visualize and quantify radiolabeled drug uptake in tumor and its dispersion throughout the tissues of the body. We have demonstrated this capability with a newly developed theranostic agent, ^123^I-MAPi. We have shown that it can access the brain when injected intrathecally and using the intratumoral osmotic pump delivery system. While the favorable pharmacokinetics was accompanied by stressful side-effects in some mice (on account of the comparatively small volumes of the murine skull), this has not been, nor is it expected to be, a limiting factor for human studies.

To conclude, we characterized the first Auger-emitting PARP inhibitor *in vivo* which presented promising therapeutic results in a pre-clinical glioma model. The physical properties of Auger emission, paired with the biological distribution of a PARP inhibitor, make it possible to speculate that a dose escalation in patients could achieve high tumoricidal doses with limited normal tissue toxicity. The 159 keV gamma ray emitted is at the sweet spot for gamma camera imaging in patients and will allow for monitoring of treatment delivery across the BBB, redistribution, and precise dose evaluation, leading to truly personalized treatment plans (fig. S5B). ^123^I-MAPi has the potential to be a new and potent clinical agent for the treatment of brain tumors when used in conjunction with new intrathecal/CED administration methods.

## Materials and Methods

### General

Sodium [^123^I]iodide in 0.1 N NaOH with a specific activity of 7.14×10^7^ GBq/g was purchased from Nordion (Ottawa, Canada). 4-(4-fluoro-3-(4-(3-iodobenzoyl)piperazine-1-carbonyl)benzyl)phthalazin-1(2H)-one was synthesized as described previously (*39*). Olaparib (AZD2281) was purchased from LC Laboratories (Woburn, MA). PARPi-FL was synthesized as previously described (*14, 40*). Water (>18.2 MΩcm^−1^ at 25 °C) was obtained from an Alpha-Q Ultrapure water system (Millipore, Bedford, MA) and acetonitrile (AcN) as well as ethanol (EtOH) were of HPLC grade purity. Sterile 0.9% saline solution (Hospira, Lake Forest, IL) was used for all *in vivo* injections. High performance liquid chromatography (HPLC) purification and analysis was performed on a Shimadzu UFLC HPLC system equipped with a DGU-20A degasser, an SPD-M20A UV detector, a LC-20AB pump system, and a CBM-20A Communication BUS module. A LabLogic Scan-RAM radio-TLC/HPLC-detector was used to detect activity. HPLC solvents (Buffer A: Water, Buffer B: AcN) were filtered before use. Purification of 2,5-dioxopyrrolidin-1-yl 3-(iodo-*^123^I*)benzoate was performed with Method 1 (flow rate: 1 mL/min; gradient: 15 min 5-95% B; 17 min 100% B; 20 min 100%-5% B); QC analysis was performed with Method 1. Method 1 was performed on a reversed phase C18 Waters Atlantis T3 column (C18-RP, 5 μm, 6 mm, 250 mm). Purification of the final product was performed on a C6 Waters Spherisorb Column (C6, 5 μm, 4.6 mm × 250 mm) with Method 2 (flow rate: 1.5 mL/min; isocratic: 0-30 min 35% B.

### Cell Culture

The human glioblastoma cell line U251 was kindly provided by the laboratory of Dr. Blasberg (MSK, New York, NY). Cells were grown in Eagle’s Minimal Essential Medium (MEM), 10% (vol/vol) heat inactivated fetal bovine serum, 100 IU2 penicillin, and 100 μg/ml streptomycin, purchased from the culture media preparation facility at MSK (New York, NY). TS543 cells are a patient-derived glioblastoma stem line kindly provided by the laboratory of Dr. Mellinghoff (MSK, New York, NY). These cells were grown in suspension in NeuroCultTM NS-A Proliferation Kit with proliferation supplement (StemCell Technologies, Cat 05751), 20 ng/mL Recombinant Human Epidermal Growth Factor (EGF) (StemCell Technologies, Cat 02633), 10 ng/mL Recombinant Human Basic Fibroblast Growth Factor (Bfgf) (StemCell Technologies, Cat 02634), 2 μg/mL Heparin (StemCell Technologies, Cat 07980), 1× antibiotic-antimicotic (Life Technologies Gibco, Cat 15240-062), 2.5 mg/mL Plasmocin (InvivoGen, Cat ant-mpp). All cells were tested for mycoplasma contamination.

### Immunoblotting

Cells were lysed in RIPA (Thermofisher Scientific, cat #89900) buffer containing protease inhibitor at 4 °C. Lysates were run on an SDS-Page gel (BioRad). Bound antibodies were detected by developing film from nitrocellulose membranes exposed to chemiluminescence reagent (#34077, Thermo Scientific). PARP1 primary antibody (Santa Cruz #sc-7150, 0.2 μg/mL) and goat anti-rabbit IgG-HRP secondary antibody (1:10000 dilution, sc-2004, SantaCruz). An anti-β-actin antibody (Sigma #A3854, 1:1000) was used as loading control.

### Synthesis of ^123^I-MAPi

^123^I-MAPi was obtained similar to synthetic procedures reported before (*26*). Firstly, 2,5-dioxopyrrolidin-1-yl 3-(iodo-^123^*I*)benzoate was obtained by adding N-succinimidyl-3- (tributylstannyl) benzoate (250 μg, 0.5 μmol) in 10 μL of AcN to a solution containing methanol (40 μL), chloramine T (9 μg, 32 nmol), acetic acid (3 μL) and ^123^I-NaI in NaOH 0.1 M (2.5 mCi). After the reaction solution was driven for 20 min at room temperature, the reaction was purified by HLPC (Method 1), and 2,5-dioxopyrrolidin-1-yl 3-(iodo-^123^*I*)benzoate at 15.1 min. The collected purified fraction was concentrated to dryness in vacuum, reconstituted in a solution of 80 μL AcN and added to a solution of 4-(4-fluoro-3-(piperazine-1-carbonyl)benzyl)phthalazin-1(2H)-one (0.3 mg, 0.9 μmol) in 20 μL DMSO, N,N,N’,N’-Tetramethyl-O-(1H-benzotriazol-1-yl)uronium hexafluorophosphate (HBTU) (0.3 mg, 0.8 μmol) in 20 μL DMSO, 4-Dimethylaminopyridine (DMAP) (0.1 mg, 0.8 μmol) in 20 μL DMSO and 10 μL triethylamine. The reaction mixture stirred in an Eppendorf ThermoMixer^®^ for 2 h at 65 °C (500 rpm). Afterwards, the reaction mixture was injected and purified by HPLC (Method 2) and ^123^I-MAPi was collected at RT = 25.5 min (RY > 70%; RP > 99%) and concentrated to dryness under vacuum. ^123^I-MAPi was formulated with 30% PEG300 / 70% saline (0.9% NaCl) for both *in vitro* and *in vivo* assays. Co-elution with nonradioactive ^127^I-PARPi reference compound confirmed the identity of the radiotherapeutic. ^123^I-MAPi was synthesized with molar activity of 3.93 ± 0.10 GBq/μmol.

### Internalization assay

Uptake of ^123^I-MAPi was tested *in vitro* (3 replicates). 5×10^5^ U251 cells were plated 24h prior to the experiment (n = 3). Media was changed and 1 h later 3.7 kBq of ^123^I-MAPi were added to the cells. For blocking, cells were incubated with a 100-fold molar excess of Olaparib 1 h before adding ^123^I-MAPi. Media was removed, and cells were washed with PBS and lysed (1 M NaOH) at different time points. The lysate was collected, and uptake was determined by radioactivity on a gamma counter.

### Viability assay

U251 GBM cells were incubated with 0 – 296 kBq of ^123^I-MAPi or equivalent dose of Olaparib overnight and then washed and incubated for 4 d in normal media (3 replicates, n = 3 each). Viability was determined by AlamarBlue assay as indicated by the manufacturer.

### Colony formation assay

Colony formation assay (CFA) was performed using U251 cells as previously described (*41*) (n = 3). Colony formation was measured at two weeks post ^123^I-MAPi treatment and compared to ^123^I-NaI. Cells were treated adding 0 – 296 kBq of compound in each well. Colony count was normalized on plating efficiency at 0 kBq for each treatment.

### Animal Work

Female athymic nude CrTac:NCr-Fo mice were purchased from Taconic Laboratories (Hudson, NY) at age 6-8 weeks. During subcutaneous injections, mice were anesthetized using 2% isoflurane gas in 2 L/min medical air. 1×10^6^ cells were injected in the right shoulder subcutaneously in 150 μL volume of 50% media/matrigel (BD Biosciences, San Jose, CA) and allowed to grow for approximately two weeks until the tumors reached about 8 mm in diameter (100 ± 8 mm^3^).

For intravenous injections, the lateral tail vein was used. Mice were warmed with a heat lamp, placed in a restrainer, and the tail was sterilized with alcohol pads before injection.

For orthotopic injections, TS543 cells (5×10^5^ cells in 2 μL growth media) were injected 2 mm lateral and 1 mm posterior to the bregma using a Stoelting Digital New Standard Stereotactic Device and a 5 μL Hamilton syringe and allowed to grow for three weeks before treatment. All mouse experiments were performed in accordance with protocols approved by the Institutional Animal Care and Use Committee of MSK and followed National Institutes of Health (NIH) guidelines for animal welfare.

### SPECT/CT imaging

SPECT/CT scans were performed using a small-animal NanoSPECT/CT from Mediso Medical Imaging Systems (Budapest, HU). For subcutaneous TS543 xenografts, ^123^I-MAPi (2.3 ± 0.5 MBq in 20 μL 30% PEG300 in 0.9% sterile saline) was administered intratumorally as single injection. At chosen time points post injection, the mice were anesthetized with 1.5-2.0%isoflurane (Baxter Healthcare) at 2 mL/min in oxygen and SPECT/CT data was acquired for 60 min.

For orthotopic TS543 tumor-bearing mice ^123^I-MAPi (614.2 kBq in 5 μL 30% PEG300 in 0.9% sterile saline) was injected intracranially using the same coordinates as for tumor cell injection (2 mm lateral and 1 mm posterior to the bregma using a Stoelting Digital New Standard Stereotactic Device and a 5 μL Hamilton syringe). At chosen time points post injection, the mice were anesthetized with 1.5-2.0% isoflurane (Baxter Healthcare) at 2 mL/min in oxygen. SPECT/CT data was collected for 60 min.

### Ex-vivo biodistribution and dosimetry

Biodistribution studies were performed in subcutaneous TS543 xenograft-bearing mice. Mice were randomized and divided in two groups (blocked and unblocked, ntotal = 6) and ^123^I-MAPi was administered intratumorally (average injected activity 1.702 ± 0.629 MBq in 20 μL, 30% PEG300 in 0.9% sterile saline). The blocked group was pre-injected (1 mg/mouse in 100 μL 30% PEG300 / 70% NaCl 0.9%) 60 min prior to treatment with Olaparib (100mM stock in DMSO). Mice were sacrificed by CO_2_ asphyxiation at 18 h post injection and counted in a WIZARD2 automatic γ-counter (PerkinElmer, Boston, MA). Uptake was expressed as a percentage of injected dose per gram (% ID/g) using the following formula: [(activity in the target organ/grams of tissue)/injected dose].

### Dosimetry

Dosimetry calculations are described in detail in the supplemental material.

### Survival studies

Mice were inoculated orthotopically at Week 0 with TS543 cells (5×10^5^ cells in 2 μL growth media). Cells were injected 2 mm lateral and 1 mm posterior to the bregma using a Stoelting Digital New Standard Stereotactic Device and a 5 μL Hamilton syringe and allowed to grow for three weeks before treatment. At Week 3 mice were intratumorally injected with ^123^I-MAPi or 30% PEG300 in 0.9% sterile saline vehicle using the same stereotactic coordinates as for tumor implantation. Mice were monitored daily thereafter. Study endpoint was determined based on animals’ sign of discomfort, pain, or significant weight loss.

### Orthotopic CED model

ALZET^®^ Osmotic Pumps were implanted subcutaneously into the mice to slowly deliver an infusion into the brain using the same coordinates as for the tumor cell injection as previously described (*26*). Control mice received an ALZET^®^ Osmotic Pump Model 1003D with Brain Infusion Kit 3 containing 30% PEG/PBS vehicle. Treatment mice received ^123^I-MAPi (average pump activity 481 ± 111 kBq in 100 μL, 30% PEG300 in 0.9% sterile saline. Delivery flow 1 μL/h, over 5 d). Pumps were surgically removed 5 d post implantation and mice monitored daily thereafter.

### Intrathecal injection

Brain-tumor-bearing mice were anesthetized and injected in the intrathecal space (injection site from L3/5 or L4/5 to prevent spinal cord injury) 2.0 MBq of ^123^I-MAPi in 50 μL 30% PEG300 in 0.9% sterile saline ^123^I-MAPi were injected and the mice imaged 1 h post injection.

### Hematoxylin–Eosin (H&E) staining

Brains were collected from mice at the time of death. The collected brains were embedded in Tissue-Tek O.C.T. compound (Sakura Finetek, Torrance, CA), flash-frozen in liquid nitrogen and cut into 10 μm sections using a Vibratome UltraPro 5000 Cryostat (Vibratome, St. Louis, MO). Sections were subsequently subjected to H&E staining for morphological evaluation of tissue pathology.

## Acknowledgements

We thank Dr. Cameron Brennan and the Brain Tumor Center for the GBM cell line model. We thank Dr. Ingo Mellinghoff and Dr. Beatriz Salinas for their helpful discussions and for providing their expertise. We thank Dr. Pat Zanzonico and Valerie Longo for technical support and help with animal imaging. We thank the Molecular Cytology Core Facility, the Animal Imaging Core Facility and the Radiochemistry & Molecular Imaging Probes Core Facility at Memorial Sloan Kettering Cancer Center. We also thank Garon Scott for editing the manuscript.

## Author contributions

G.P., S.A.J. and T.R. conceived the study and designed the experiments. G.P., S.A.J., A.S., N.G., P.D.S.F., S.K., and M.M. carried out experiments and collected data. G.P., S.A.J., L.C., B.M.Z., J.S.L. J.L.H. and T.R. analyzed the data. G.P., S.A.J., L.C. J.L.H. and T.R. wrote the manuscript. All authors carefully reviewed and approved the manuscript.

## Data and materials availability

All data necessary for interpreting the manuscript have been included in the main manuscript or in the Supplementary Materials. Additional information may be requested from the authors.

