## Supplemental material for "Targeted brain tumor radiotherapy using an Auger emitter"

### Supplemental material table of contents

**Dosimetry (organ-level/3D) of $^{123}$I-MAPi via CED administration**

*Estimation of time-integrated activity coefficients in mouse organs*

Radiation dose estimates for osmotic pump-based intratumoral administration of $^{123}$I-MAPi were obtained under the assumption that biological clearance kinetics of a small impulse of $^{123}$I-MAPi are equivalent to those of $^{131}$I-PARPi administered as a single bolus intratumoral injection, and further that the clearance kinetics are not concentration-dependent. Under these assumptions the activity $A(r_S, t)$ (MBq) of $^{123}$I-MAPi in organ S is given by:

$$A(r_S, t) = \dot{A}(t) * a'(r_S, t) \qquad \text{Eqn. 1}$$

where $*$ is the convolution operator, $\dot{A}(t)$ is the rate of administration (MBq/h), and $a'(r_S, t)$ is the time-dependent fraction of injected activity of an impulse of $^{123}$I-MAPi in organ S (i.e. that of a single bolus). In the case that the dose integration period $T_D$ is taken as infinity (the usual case), the time-integrated activity coefficient for the present case (slow infusion given by $\dot{A}(t)$) is equivalent to the case of a single bolus injection. Thus, time-integrated activity coefficients for mouse organs were calculated by applying the appropriate decay factor for $^{123}$I to previously-obtained organ time-activity data *(1)* for intratumorally-injected $^{131}$I-PARPi, and performing trapezoidal integration of the adjusted data for each organ (Table S1). Clearance after the last measured time point was assumed to occur via radioactive decay only.

*Estimation of time-integrated activity coefficient for osmotic pump contents*

Under the assumption that the $^{123}$I-MAPi is homogeneously distributed within the osmotic pump contents, which occupy an initial volume $V_0$ (µL) and are injected at a constant volumetric flow rate $\kappa$ (µL/h), the activity of the pump contents $A(pump, t)$ can be expressed as (*1*):

$$A(pump, t) = H\left(\frac{V_0}{\kappa} - t\right) \cdot A_0\left(1 - \frac{\kappa t}{V_0}\right) e^{-\lambda t} \qquad \text{Eqn. 2}$$

where $t$ is the time post-implantation, $A_0$ is the initial activity in the implanted pump, and $H(V_0/k - t)$ is a Heaviside function shifted in time by $V_0/k$ (i.e. the time the pump becomes empty).

The activity injected $A_{0,inj}(t)$ can be expressed as:

$$A_{0,inj}(t) = H(V_0/\kappa - t) \cdot \frac{\kappa A_0}{\lambda V_0}\left(1 - e^{-\lambda t}\right) + H(t - V_0/\kappa) \cdot \frac{\kappa A_0}{\lambda V_0}\left(1 - e^{-\lambda V_0/\kappa}\right) \qquad \text{Eqn. 3}$$

We define the time-integrated activity coefficient for the pump contents as normalized to the activity injected (i.e. delivered through the cannula):

$$\tilde{a}(pump, t) = \frac{\int_0^t A(pump, t)dt}{A_{inj}(t)} \qquad \text{Eqn. 4}$$

which for $t$ = $T_D \rightarrow \infty$ is:

$$\tilde{a}(pump) = \frac{\lambda\kappa - \kappa + \lambda V_0}{\lambda\kappa\left(e^{\lambda V_0/\kappa} - 1\right)} - \frac{\kappa - \lambda V_0}{\lambda\kappa} \qquad \text{Eqn. 5}$$

The manufacturer specified values for $\kappa$ (1 µL/h) and $V_0$ (100 µL) were used for estimation of $\tilde{a}(pump)$ (fig. S5A, S5B, S5C, Table S1).

**Supplementary Table 1:** Time-integrated activity coefficients estimated for intratumoral administration $^{123}$I-MAPi via osmotic pump.

| Organ or Region | Time-integrated activity coefficient (MBq*h/MBq) |
|---|---|
| Intestines | 0.399 |
| Liver | 0.208 |
| Pancreas | 0.000886 |
| Stomach contents | 0.0117 |
| Heart contents | 0.00536 |
| Heart wall | 0.00527 |
| Lungs | 0.00640 |
| Brain | 0.000946 |
| Thyroid | 0.000699 |
| Skeleton | 0.0597 |
| Tumor | 0.0597 |
| Kidneys | 0.00617 |
| Remaining organs | 0.139 |
| $^{123}$I-MAPi solution (pump contents) | 81.5 |
| Osmotic pump (container portion) | 0 |

*Construction of a CT-derived phantom for dose calculation*

A computed tomography (CT) scan of a mouse bearing an osmotic pump implanted over the left shoulder was acquired on a NanoSPECT/CT device from Mediso Medical Imaging Systems (fig. S5C). CT scans were reconstructed with an isotropic voxel size of 0.147 × 0.147 × 0.147 mm using a Feldkamp cone-beam filtered back-projection algorithm (Mediso Ltd.). A combination of manual and semi-automatic segmentation techniques were used to define the osmotic pump, tumor, and major organs within 3D Slicer 4.10 (www.slicer.org); the segments were subsequently converted to triangulated polygonal mesh. Within the mesh-editing software Blender (www.blender.org), the polygon count of each mesh was reduced using an edge-collapse decimation modifier (ratio 0.05-0.10), and small modifications made to correct for any overlapping/non-manifold geometry. Thin wall structures of select organs (e.g. stomach, bladder) were modeled by applying a vertex offset modifier to the corresponding structure defined previously via segmentation. Finally, all surface meshes were merged, exported, and converted to a tetrahedral mesh using TetGen *(2)*. Following definition of the phantom region information (region name, composition, etc.) using an in-house python program, the completed phantom was imported into the PHITS-Based Application for Radionuclide Dosimetry in Meshes (PARaDIM v1.0).

*Absorbed dose calculation in PARaDIM*

Absorbed doses were calculated with PARaDIM v1.0 using time-integrated activity coefficients given in Table S1. $1\times10^8$ total histories were simulated via PHITS on a HP Z8 workstation (3.6 GHz Intel Xeon 5122 processor, Windows 10 OS, single core calculation). The default settings for physical models and energy cutoff (1 keV for electrons and photons) were used. The material surrounding the phantom was defined as void. An isotropic voxel resolution of 1.0 mm was used for 3D dose calculation.

### Supplemental figures

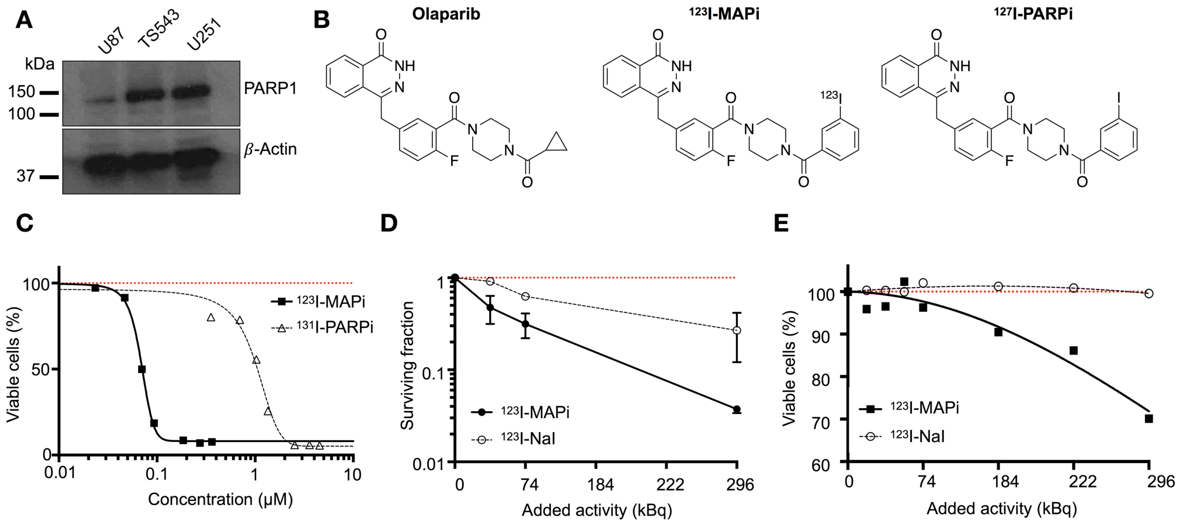

**Supplemental Figure 1. Molecular structures, target expression, clonogenic assay and viability assay *in vitro*. A.** Western Blot of PARP1 in glioblastoma cell lines with $\beta$-actin control. **B.** Molecular structure of Olaparib, $^{123}$I-MAPi, and $^{127}$I-PARPi, $^{123}$I-MAPi isotopologue. **C.** Alamar Blue assay comparing $^{123}$I-MAPi and $^{131}$I-PARPi at equal molar concentrations. $^{123}$I-MAPi $EC_{50}$ = 69 nM, $^{131}$I-PARPi $EC_{50}$ = 1148 nM. Nonlinear fit four parameters variable slope. **D.** Clonogenic assay and **E.** Almar Blue viability assay of $^{123}$I-MAPi vs. sodium iodine-123 in U251 GBM cells.

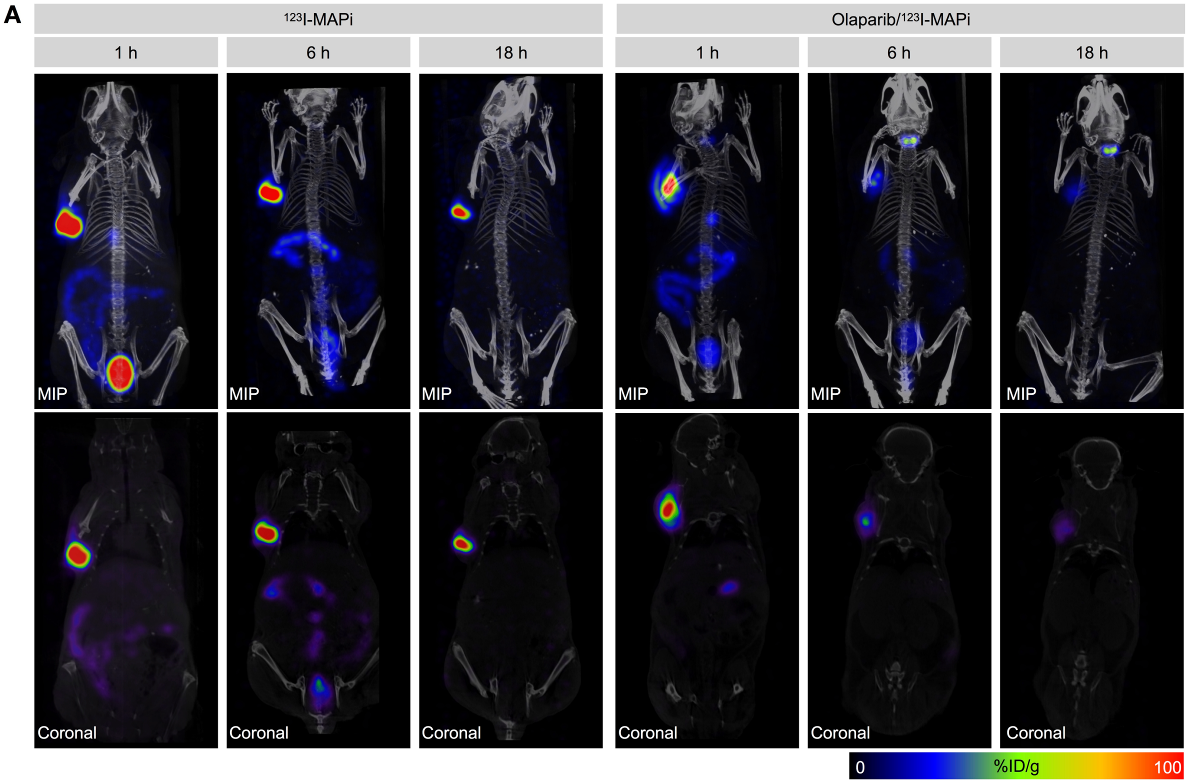

**Supplemental Figure 2. *In vivo* SPECT-CT of $^{123}$I-MAPi in Subcutaneous TS543 model.**

**A.** MIPS (top row) and corresponding quantified coronal slices (bottom row) at selected time points after intratumoral injection of $^{123}$I-MAPi with or without intravenous blocking with Olaparib.

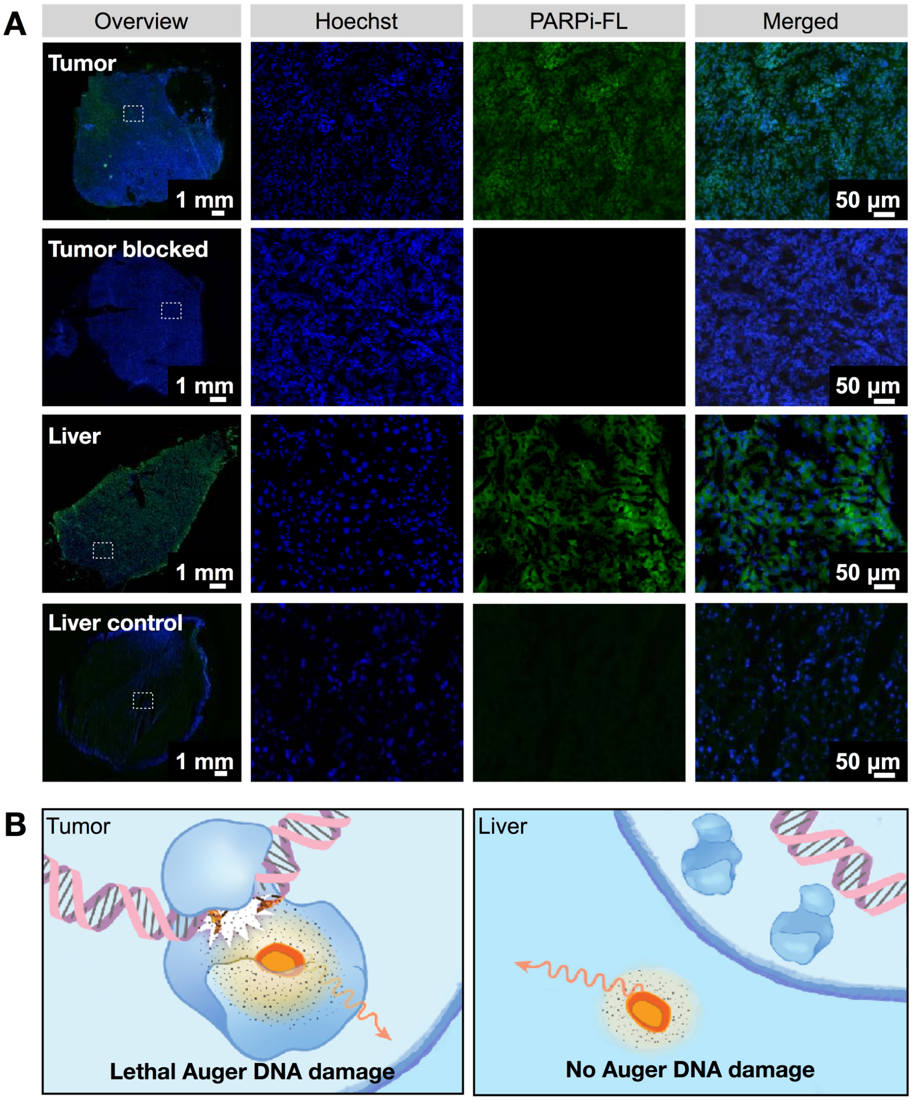

**Supplemental Figure 3. Subcellular localization study for $^{123}$I-MAPi. A.** Tissue from mice injected with PARPi-FL and Hoechst. PARPi-FL uptake was blocked in tumors by injection of 100-fold excess dose Olaparib 1 h prior to injection of PARPi-FL. Liver control was performed injecting vehicle IV. **B.** A suggested mechanism of radiation protection for liver cells is that Auger radiation specific toxicity depends on the isotope being in the proximity of the target DNA, which doesn't happen in these cells.

A

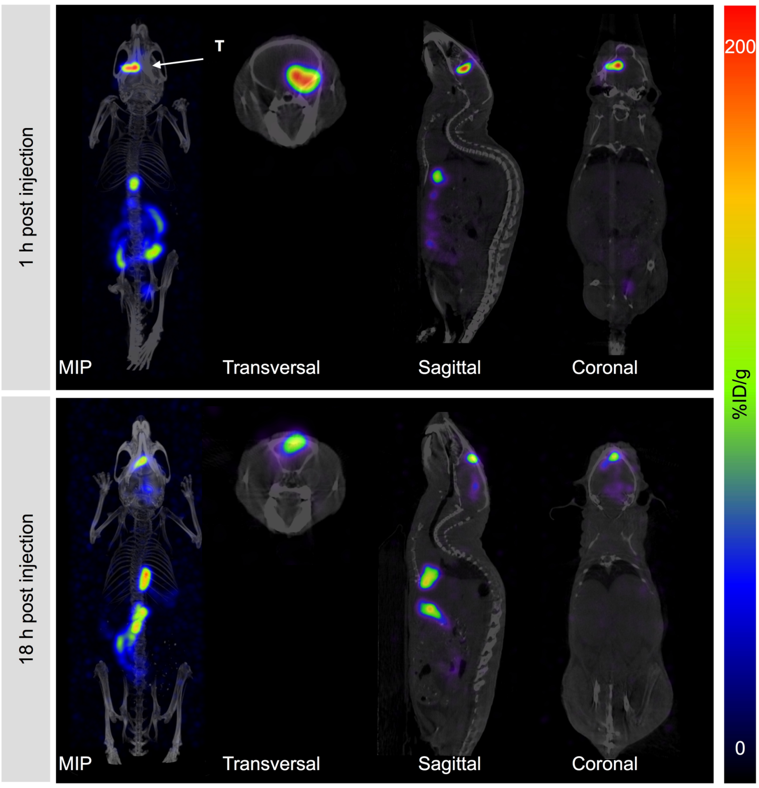

**Supplemental Figure 4.** ***In vivo*** **SPECT-CT of** $^{123}$**I-MAPi in orthotopic TS543 model. A.** Quantified coronal slices at 1 h and 18 h post local injection of $^{123}$I-MAPi. $^{123}$I-MAPi was injected intratumorally using the same stereotactic coordinates as for tumor implantation. The animal was imaged in a nanoSPECT/CT at the indicated time points post injection.

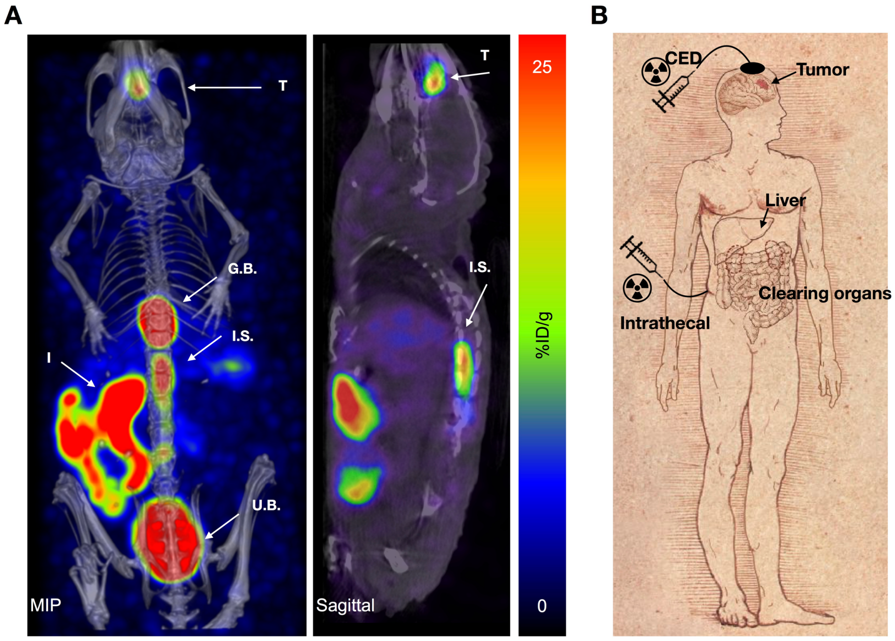

**Supplemental Figure 5. Translation potential of $^{123}$I-MAPi. A.** SPECT/CT imaging of $^{123}$I-MAPi administered intrathecally, 1 h post injection. **T**: tumor; **I.S.**: injection site; **G.B.**: gallbladder; **I**: intestine; **U.B.**: urinary bladder. **B.** We speculate that the different delivery methods characterized can show clinical potential for a reduced patient's stress and improved tumor accumulation. Biodistribution of $^{123}$I-MAPi in the cytosol of non-target cells limits unspecific toxicity and clinical side effects, preserving cleaning organs.

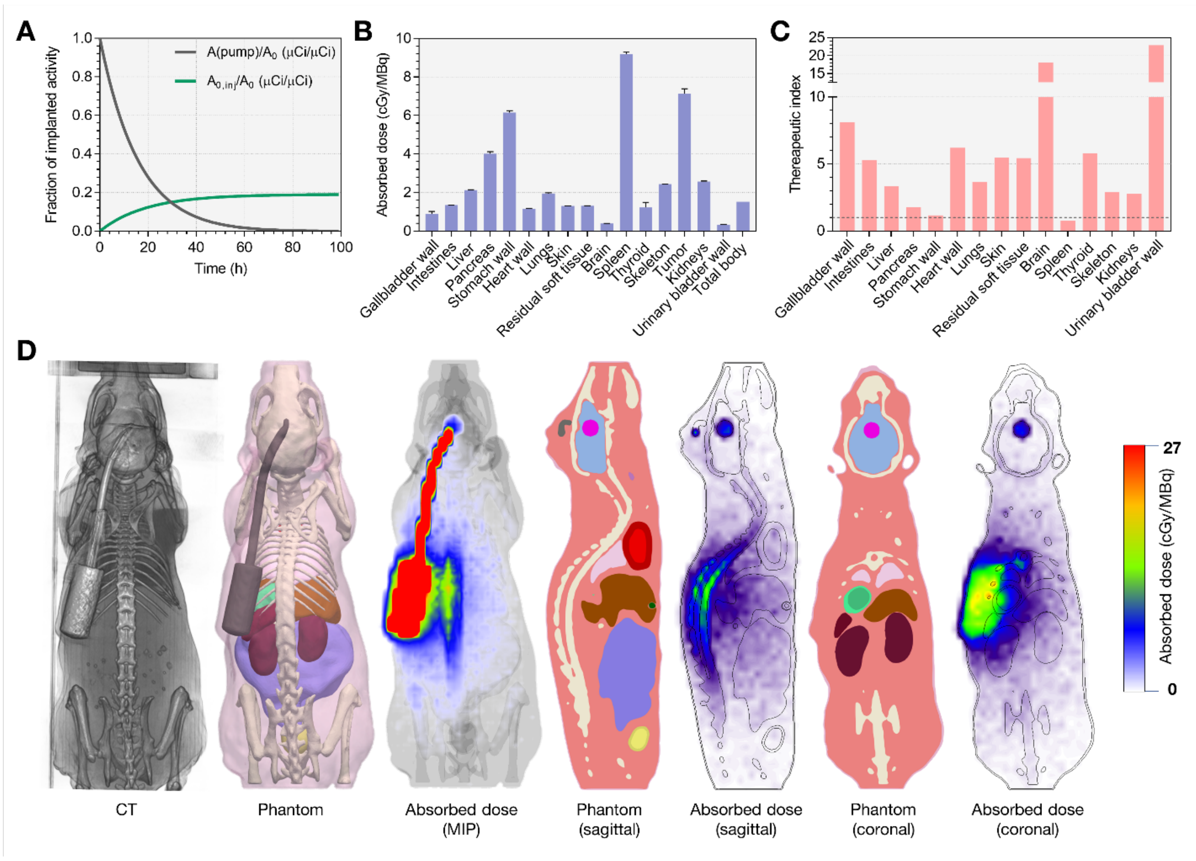

**Supplemental Figure 6. Radiation dose estimation for intratumoral administration of $^{123}$I-MAPi via CED-mimicking osmotic pump. A.** Time-dependent activity of pump contents (gray), and injected activity (green), normalized to the initial activity $A_0$ of the pump contents. **B.** Mean absorbed doses for whole organs computed in CT-derived phantom shown in (C). Error bars represent statistical uncertainties in Monte Carlo dose calculation. **C.** Therapeutic indices calculated as ratio of mean tumor dose to mean normal organ doses. **D.** 3D absorbed dose calculation demonstrates elevated dose in regions in proximity to activity reservoir of pump.

**Supplemental material references**

1. S. A. Jannetti *et al.*, *J Nucl Med* **59**, 1225 (2018).
2. H. Si., *ACM Trans Math Softw* **41**, 1 (2015).
